# Mapping Chromatin Interactions of ZBP1 and ADAR1 Z-Alpha Domains: A ChIP-Seq Based Comparison

**DOI:** 10.1101/2024.11.29.626086

**Authors:** Dennis Hamrick, Manjita Sharma, Edward Grow

## Abstract

The DNA double helix typically exists in the canonical B-form conformation, but this structure often can adopt the unique alternative form known as Z-DNA. In Z-DNA, the DNA helix winds to the left in a zigzag pattern instead of the right-handed B-DNA form. Z-DNA is thought to play a key role in transcription, but it is unclear whether is a positive or negative regulator of RNA polymerase activity. Additionally, several studies have shown how Z-DNA contributes to DNA damage or genome instability. However, the precise role of Z-DNA in the genome remains unclear. To address this question, we mapped Z-DNA using a ChIP-Seq assay with two Z-DNA biosensors: Zaa-Zbp1, comprised of a dimerized Z-alpha Z-DNA binding domains from Z-DNA binding protein 1 (Zbp1), and Zaa-Adar1, comprised of dimerized Z-alpha domains from Adenosine deaminase acting on RNA 1 (Adar1). We found that these Zaa probes possessed similar binding profiles when analyzed with motif analysis, but gene ontology analysis revealed that these Z-alpha domains bound to heterogeneous genes, with Zaa-Zbp1 most strongly binding to genes in the RHOQ-GTPase pathway and Zaa-Adar1 binding to genes involved in the M phase of the cell cycle.

## 1 Introduction

### 1.1 Z-DNA History

Z-DNA was accidentally discovered in 1979, when researchers performing single-crystal X-ray diffraction on a d(CG)3 DNA hexamer revealed a dramatically different structure from the expected smooth right-handed double helix. Instead, the experiment revealed a left-handed double helix with alternating syn and anti-conformation bases, as well as a zig-zagging phosphate backbone that earned it the name Z-DNA[1]. Z-DNA was shown to be more likely to form in high salt conditions, in purine-pyrimidine repeat sequences, and was associated with negative supercoiling of the DNA molecule. dsRNA was found to also be capable of adopting a Z-form under similar conditions [2]. After an initial surge of interest, Z-DNA and Z-RNA were largely sidelined as a mild curiosity without biological relevance.

Work in the following years demonstrated that Z-DNA formed *in vivo* and could be mapped with Z-DNA specific antibodies. Computational analysis of 137 genes in the human genome revealed that Z-DNA forming sequences were enriched near transcription start sites[3]. Experimental work showed that Z-DNA formed at the transcription start site for the gene C-MYC when it was being actively transcribed, and this formation was ceased by halting C-MYC transcription [4]. This resulted in the model that Z-DNA is formed by negative supercoiling behind active RNA polymerases, and the cell uses topoisomerase to relax this negative supercoiling and reverse the formation of Z-DNA [1].

In 1995, a major breakthrough in determining Z-DNA’s biological relevance came when Herbert et al. showed that the dsRNA-editing protein ADAR1 possesses a Z-nucleic acid binding domain, christened Z*α*. Z*α* domains were subsequently found in a myriad of other proteins [5]. The Z*α* containing proteins experimentally characterized thus far are either associated with the innate immune response and probably largely bind to cytoplasmic Z-RNA (ADAR1, ZBP1, PKZ) or are viral homologues of these defensive proteins that counteract the host innate immune response (E3L, ORF112).

### 1.2 Z*α* Domains

Z*α* domains are Z-DNA-specific binding domains that bind to the sugar-phosphate backbone. They are non-sequence specific and can bind it and stabilize the Z-formation. We have mainly focused on ADAR1 and ZBP1. The adenosine deaminase (ADAR) is specifically involved in immune response and ZBP1 which is a DNA-binding protein. Before the Z*α* domain was discovered, there were several non-physiological studies conducted in Z-DNA. The discovery of Z*α* domain is the most specific biosensor of Z-DNA and was used for Chip-Seq where they were expressed in cells and were bound to Z-DNA.

ADAR1 (Adenosine Deaminase Acting on RNA) and ZBP1 (Z-DNA Binding Protein 1) play a distinct role in immune response regulation and cellular biology. The Z*α* domain or the Z-DNA binding domain, that allows interaction with Z-DNA, is found in both proteins. On the other hand, ZBP1 is a specific protein that binds to Z-DNA and plays a pivotal role in cellular response. The Z*α* domain within the ZBP1 protein activates immune responses that help fight infections.

ADAR1 possesses two isoforms, p150 and p110. Both isoforms contain catalytic A-to-I editing domains and a dsRNA binding domain, with only the p150 isoform also containing an N-terminal Z*α* domain. This Z*α* domain is coupled with a Z-beta domain that does not bind Z-DNA and has an unclear function. This Z*α* domain functions to recognize cytoplasmic dsRNA in the Z conformation, especially Long Interspersed Nuclear Element-1 (LINE-1) and Alu element RNAs. This recognition enables ADAR1 to edit adenosines to inosine in these elements, blocking recognition by antiviral sensors like RIG-I and preventing a strong innate immune response [6]).

While ADAR1 serves to dampen the immune response to self-generated Z-RNA’s, ZBP1 serves the opposite function. ZBP1 is upregulated by IFN-1 in response to viral infection. ZBP1 binds cytoplasmic Z-RNAs generated by viral infection and initiates a PANoptotic response by activating the NLRP3 inflammasome, culminating in PANoptotic cell death. ZBP1 is also similarly implicated in necroptosis and pyroptosis [7].

Both ADAR1 and ZBP1 stand out as the only Z-nucleic acid sensing proteins identified in mammals thus far, with complementary roles in the innate immune response. Previously, a Z-DNA binding construct has been created by excising the N-Terminus region of ADAR1p150 and replacing the Z-Beta domain with another Z*α* domain. This construct, referred to as Z*αα*, possesses a significantly increased affinity for Z-DNA compared to the wild-type N-terminus of ADAR1. This ADAR1-Z*αα* construct was used for ChIP-SEQ in HeLa cells, which found that Z-DNA was associated with actively transcribed genomic regions [8].

We first asked the question: where in the genome is Z-DNA located? To address this question, we performed two ChIP-SEQ experiments: one using ADAR1-Z*αα*, and the other using a novel ZBP1-Z*αα* probe. We performed these experiments in R1 mouse embryonic stem cells (mESCs) due to their pluripotent state and amenability to genetic modification. This methodology has generated the first ever genome-wide map of Z-DNA in the mouse genome, as well as the first map of the genomic binding profile of a non ADAR1 Z*αα* construct. Additionally, we have utilized the Z-HUNT algorithm and updated it for use in modern big data genomic workflows. This updated version, Z-Nome-HUNT, enables us to annotate the genome with computationally predicted sites of Z-DNA formation and compare our experimental data to these predictions.

## 2 Methods

### 2.1 Cell Culture and Transgene Expression

Following methods described in Wang et al. 2022, we created an R1 mouse embryonic stem cell (mESC) population possessing a doxycycline inducible Za2xZa (NESmutant)-HA transgene. Expression of the Z*αα* dimer was confirmed by inducing over-expression with 1 ug/ml doxycycline added to media for 18 hours, then harvesting cells with 0.01% trypsin, lysing with RIPA buffer, and performing a western blot with Abcam anti-HA tag ab9110. Immunofluorescence was also performed to confirm the presence of the transgene and localization within the nucleus. Cells were incubated in 2i-LIF media with 1ug/ml doxycycline for 18 hours, then fixed with 4% PFA for 15 minutes. They were then blocked and permeabilized with 5% BSA, 0.3% Trition in PBS, then incubated at 4 C with Abcam ab9110 anti-HA at 1/1000 dilution in 1% BSA, 0.1% Triton solution with rotation overnight. The next day, cells were washed with PBS 3x for ten minutes and stained with anti-Rabbit 594 at 1/1000 dilution and DAPI at 1/2000 dilution.

### 2.2 Chromatin Immunoprecipitation

To identify the genomic role of Z-DNA, we performed a Chromatin Immunoprecipitation experiment using these cells. Expression of Za2xZa was induced by overnight incubation of the population with 1 ug/ml doxycycline. The cells were rinsed with PBS and then were treated with 1% formaldehyde solution. After crosslinking them at room temperature for 10 minutes, they were harvested, and ChIP was performed with these crosslinked samples.

Referenced from [9], the samples were treated with the following modifications. The samples were first lysed with lysis buffer 1 (50mM HEPES, 150mM NaCl, 1mM EDTA, 10% glycerol, 0.5% NP40, 0.25% Triton). The pellets were again lysed with lysis buffer 2 (10mM Tris, 200mM NaCl, 1mM EDTA, 0.5M EGTA). Then, the pellets were sonicated in lysis buffer 3 (10mM Tris, 100mM NaCl, 1mM EDTA, 0.5 mM EGTA, 0.1% Na-Deoxycholate, 0.5% N-lauroylsarcosine) with a Covaris ultrasonicator. The sonicated samples were treated with HA-tag antibody (Rabbit, polyclonal, ab9110). The samples were then treated with RNAse and proteinase K and purified with PCI extraction. The DNA was precipitated, and the ChIP-seq libraries were then created using the NEBnext ChIP-seq library kit and sequenced with Illumnia Novoseq.

Z-Nome-HUNT was modified based off of the M-Hunt algorithm source code published on the Ho Lab GitHub[10]. Z-Nome-HUNT was used with settings max = 12, min = 6, windowsize =12 on the mm10 genome to generate Z-DNA predictions for every base in the genome, which were then averaged for every 20 bases and filtered for Z-scores greater than 700. Z-Nome-HUNT source code and binaries are available at https://github.com/dennishamrick/z-nome_hunt, and a web-based GUI application is available at http://dennis-hamrick.shinyapps.io/z-nome_hunt.

### 2.3 Data Analysis

Analysis was performed using R version 4.4.2 and the Galaxy webserver. Gene ontology analysis (Figure 3a & 3b) was performed using ReactomePA [11]. A heatmap (Figure 1b) was generated using deepTools version 3.5.4 [12] Figures 2a and 2b were generated using R package ggplot2 version 3.5.1. [13]. The eulerr R package was used for generating Figure 2e[14]. Motif analysis was carried out using GimmeMotifs version 0.18.0[15][16].

**Figure 1:**
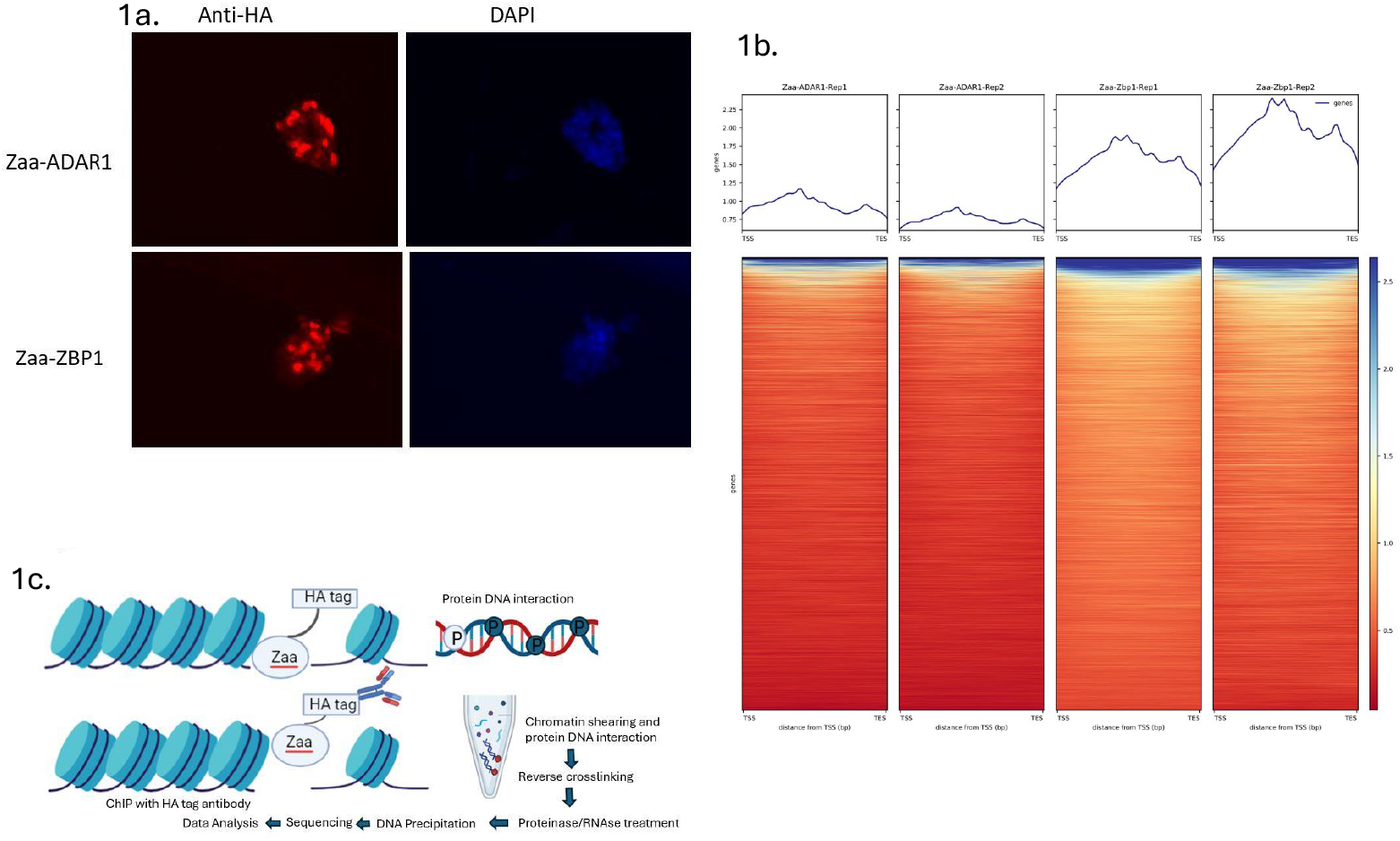
a. Immunofluorescence results for Z*αα*-Adar1 and Z*αα*-Zbp1 constructs. DAPI - 4’,6-diamidino-2-phenylindole. b. Heatmap results from deepTools on both ChIP-Seq samples. N = 8183 regions. c. Diagram of Z*αα*-ChIP-Seq process.

**Figure 2:**
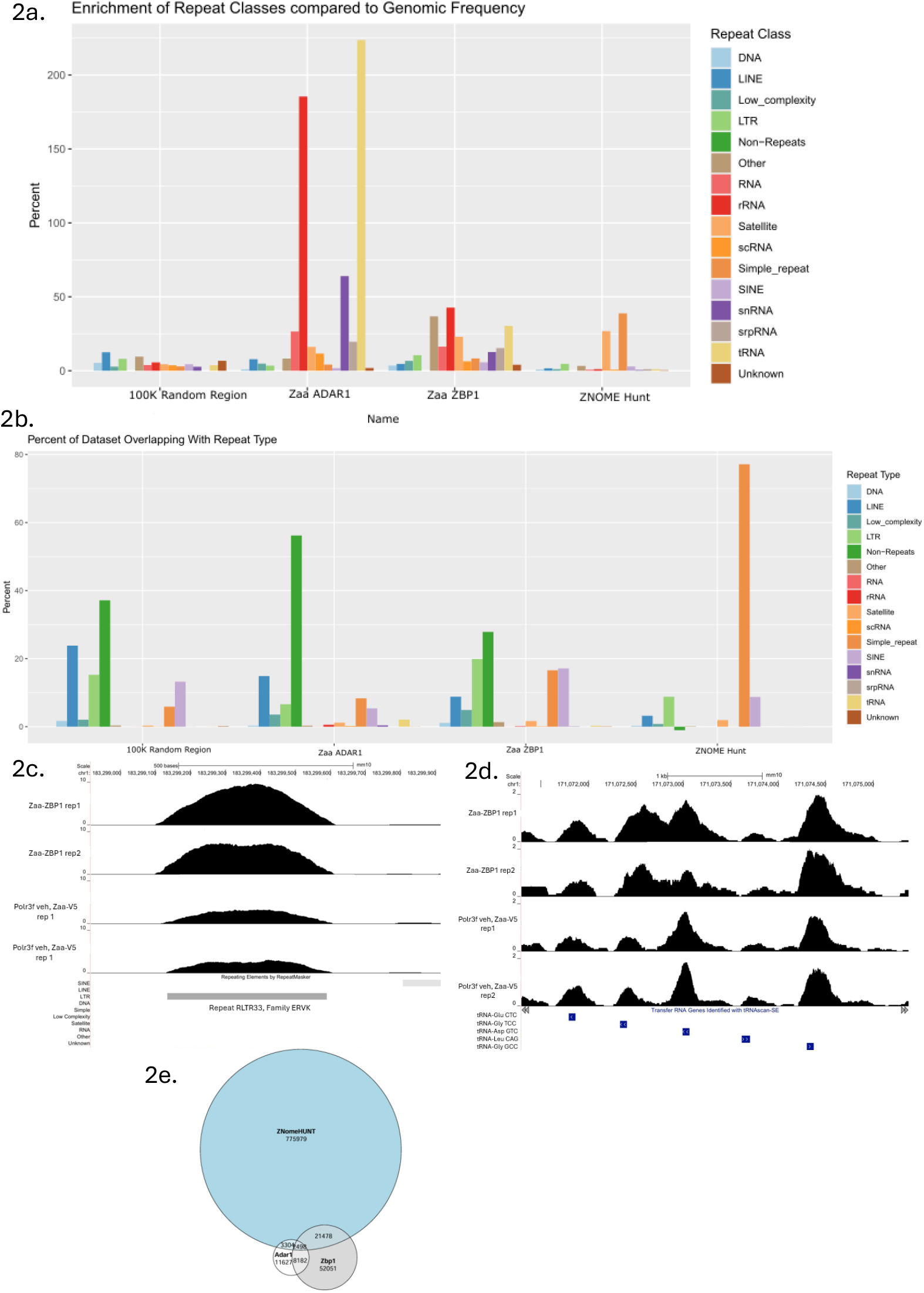
a. Enrichment of repeat classes in each dataset compared to genomic frequency. Number of peak regions for each dataset: Z*αα*-ADAR1: 11627, Z*αα*-Zbp1: 52051, Z-Nome-HUNT:775979. b. Percent of each dataset overlapping with each repeat type. c. UCSC Genome Browser image showing Z*αα*-ZBP1 peak at an ERVK repeat. d. UCSC Genome Browser image showing peaks at tRNA genes.

**Figure 3:**
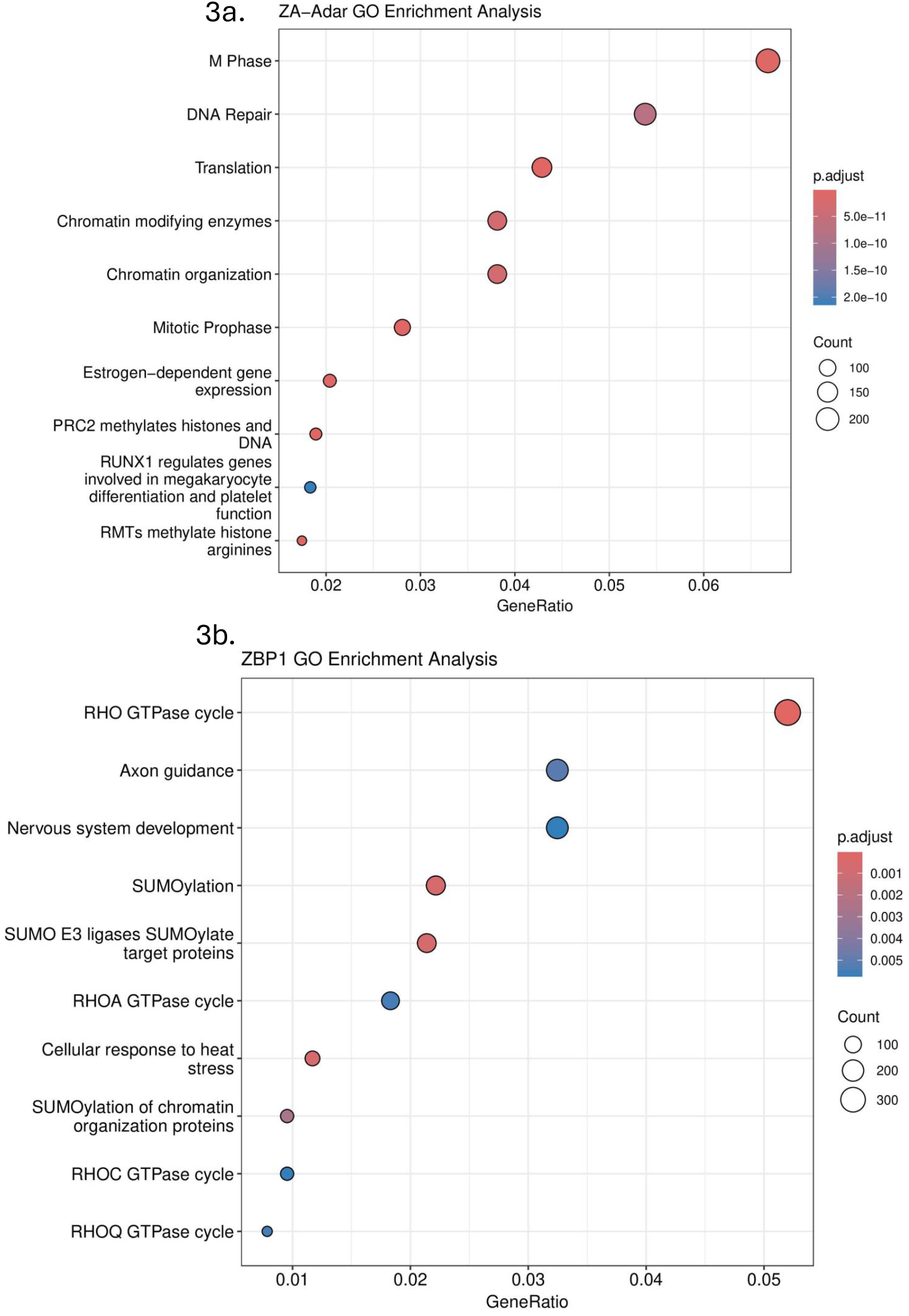
a. Results of Z*αα*-ZBP1 ReactomePA GO Enrichment Analysis. b. Results of Z*αα*-ADAR1 GO Enrichment Analysis.

## 3 Results

### 3.1 Repeat Enrichment

We first tested whether Z-DNA was localized to repeat regions of the genome. Z*αα*-ADAR1 and Z*αα*-ZBP1 had markedly different profiles of binding at various genomic repeats (Figure 2a). Both groups differed from random binding profiles, and both groups also did not match with Z-Nome-HUNT binding, indicating that sequence susceptibility to Z-DNA formation was not the major factor leading to Z-DNA formation and Z*α* binding at these sites. Most strikingly, Z*αα*-ADAR1 binding sites were highly enriched at tRNA genes (a 223% enrichment) and at rRNA sites(a 185% enrichment). This effect was present more weakly in the Z*αα*-ZBP1 data, with Z*αα*-ZBP1 also having less apparent bias for any one repeat type.

**Table 1:**
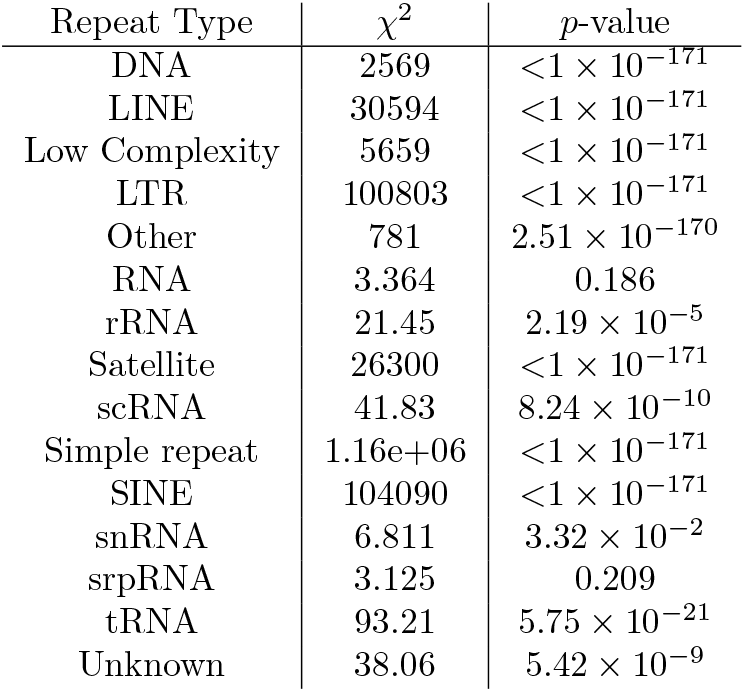
Results of *χ*^2^ test for Z*αα*-ZBP1, Z*αα*-ADAR1, and Z-Nome-Hunt. *****************************A *χ*^2^ test revealed that for all annotated RepeatMasker groups besides “RNA” and “srpRNA”, there was a statistically significant difference between the *Zαα-ADAR1*,*Zαα-ZBP1*, and Z-Nome-Hunt results.

### 3.2 GO Enrichment Analysis

#### 3.2.1 Z*αα*-ZBP1 GO Enrichment Analysis

The Z*αα*-ZBP1 Gene Ontology (GO) Enrichment Analysis plot (Figure 3a) shows different pathways and their gene ratio in which the analysis is done. The gene ratio is highest for the RHO GTPase cycle, with 5.2% of gene-localized peaks being involved with this process,. GTPases such as RHO are important for cellular processing and are controlled through their active and inactive sites. The Rho GTPase cycle is essential for cellular processes such as activation, inactivation, cell proliferation, and survival (Boureux, 2006). The GTPases regulates remodeling which is essential for innate immune responses. This innate immune response has a major involvement with ZBP1 protein-which detects Z-DNA. The cycles are regulated by Z-DNA binding proteins through immunoregulatory pathways[17].

The RHO GTPase cycle plays an essential role in regulating the cellular processes through biochemical networks. When there is DNA damage, RHO GTPase is activated, and the pathways are repaired after activation which maintains the integrity of genome through cellular responses. This pathway is also essential for regulation of actin cytoskeleton which maintains the shape of the cell and facilitates repair of DNA. Hence, this pathway of DNA repair and damage is coordinated by RHO GTPase. It responds to DNA damage by homologous and non-homologous recombination. This contributes to DNA stress and repairs damage [18].

Our Z*αα*-ZBP1 analysis shows the highest enrichment in the RHO GTPase cycle. ZBP1 protein when bound to Z-DNA is involved in cellular response to stress which can be related to the RHO GTPase that plays a crucial role in influencing cellular response through cytoskeleton and DNA damage. This remodeling could potentially affect the formation of Z-DNA from response through stress leading to the hypothesis that GTPases could influence the binding of ZBP1 to Z-DNA. This can lead to changes in cellular structures leading to Z-DNA formation [19].

#### 3.2.2 Z*αα*-ADAR1 GO Enrichment Analysis

Our GO enrichment results from Z*αα*-ADAR1 (Figure 3b) overlapped well with the GO enrichment results from the Shin et al Z*αα*-ADAR1 ChIP-Seq. Both analyses found enrichment at genes involved in translation, the regulation of the cell cycle, and chromatin organization and modification. These genes include ribosomal proteins (Rps5, Rps9), translation elongation and initiation factors (Eef2, Eif5), and DNA methyltransferases (Dmnt1, Dmnt3b). Given that only ADAR1p110, an isoform missing the Z*α* domain, is found in the nucleus, it is unlikely that ADAR1 or ZBP1 bind to nuclear Z-DNA. This is further evidenced by the low overlap of Z-Nome-HUNT predicted Z-DNA sites with Z-DNA sites mapped by either Z*αα* construct, indicating that processes beyond sequence susceptibility regulate the formation of Z-DNA.

### 3.3 Motif Analysis

Motif analysis, performed with GimmeMotifs 0.18.0, revealed classical Z-DNA forming sequences as motifs for both proteins(Figure 4). Most motif patterns recognized were variations on purine-pyrimidine repeats such as CG_*n*_ or CA_*n*_.

**Figure 4:**
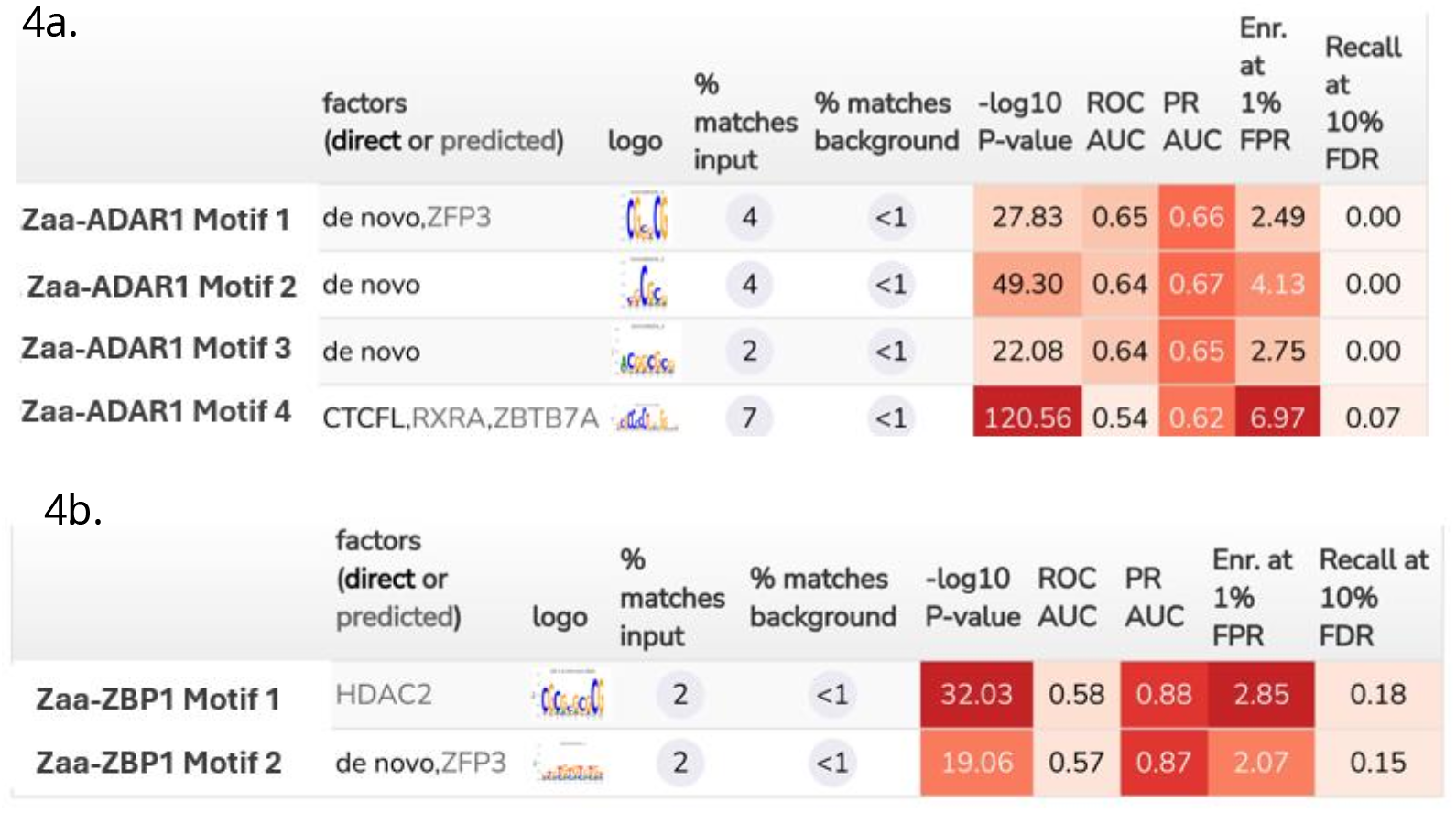
a. GimmeMotifs analysis for Z*αα*-ADAR1.b. GimmeMotifs analysis for Z*αα*-ZBP1.

## 4 Conclusion

In this study, we used ChIP-Seq to compare the binding profiles of two Z-DNA binding domains: the Za domain from ADAR1 and the Za domain from ZBP1. Our findings demonstrate that these two probes, while possessing similar structural affinities for Z-DNA, exhibit divergent binding patterns at specific genomic sites. Notably, Z*αα*-ADAR1 showed significant enrichment at genes involved in M-phase, translation, tRNA genes and rRNA genes, suggesting a potential regulatory role of Z-DNA in translation, ribosome biogenesis, and other fundamental cellular processes. In contrast, Z*αα*-ZBP1 displayed notable enrichment in genes involved in the RHO GTPase cycle, indicating that ZBP1 may play an essential role in cellular responses to stress and immune signaling through Z-DNA binding. These results expand our understanding of the genomic role of Z-DNA. Much of the work in this field thus far has been focused on elucidating the function of Z-NAs in the innate immune response, a clearly important role given the multiple pathways involved in sensing cytoplasmic Z-NAs. However, it is clear from the abundance of Z-DNA forming sites predicted with Z-Nome-HUNT and experimentally detected with Z*αα*-ChIP-Seq that this alternate form of DNA also plays a role in the nucleus. Previous research has determined that Z-DNA forming sequences are enriched at transcription start sites and form Z-DNA during transcription, such as at the TSS for C-MYC. This negative supercoiling-induced Z-DNA is relaxed by the action of topoisomerases[4]. We speculate that Z-DNA formation can be more precisely regulated by the cell via unrecognized nuclear DNA binding proteins with cryptic Z*α* like sequences.

Further research should aim to identify potential examples of these proteins. Additionally, further Z*αα*-ChIP-Seq experiments with additional Z*α* domain probes (such as the Z*α* domains of PKZ and E3L) would help determine the genomic binding specificity of Z*α* domains, as well as refine the Z-DNA profile of various cell types and specific factors influencing Z-DNA formation. Understanding the regulatory mechanisms behind Z-DNA formation and stabilization will provide deeper insights into its role in genome organization, transcription regulation, and the innate immune response. Additionally, leveraging updated algorithms like Z-Nome-HUNT alongside experimental validation will help refine our understanding of the complex interplay between alternative DNA structures and their function.

## 5 Acknowledgments

This work was supported by CPRIT grant RR210077.

## References

[1] Alexander Rich and Shuguang Zhang. “Timeline: Z-DNA: the long road to biological function”. en. In: Nat. Rev. Genet. 4.7 (July 2003), pp. 566–572.

[2] Parker J Nichols et al. “Z-RNA biology: a central role in the innate immune response?” en. In: RNA 29.3 (Mar. 2023), pp. 273–281.

[3] G. P. Schroth, P. J. Chou, and P. S. Ho. “Mapping Z-DNA in the human genome”. In: Computer-aided mapping reveals a nonrandom distribution of potential Z-DNA-forming sequences in human genes. The Journal of biological chemistry 267.17 (1992).

[4] B Wittig et al. “Transcription of human c-myc in permeabilized nuclei is associated with formation of Z-DNA in three discrete regions of the gene”. en. In: EMBO J. 11.12 (Dec. 1992), pp. 4653–4663.

[5] Alan Herbert. “Z-DNA and Z-RNA in human disease”. en. In: Commun. Biol. 2.1 (Jan. 2019), p. 7.

[6] Yuhan Zhong et al. “Zα domain proteins mediate the immune response”. en. In: Front. Immunol. 14 (Sept. 2023), p. 1241694.

[7] Rajendra Karki and Thirumala-Devi Kanneganti. “PANoptosome signaling and therapeutic implications in infection: central role for ZBP1 to activate the inflammasome and PANoptosis”. en. In: Curr. Opin. Immunol. 83.102348 (Aug. 2023), p. 102348.

[8] So-I Shin et al. “Z-DNA-forming sites identified by ChIP-Seq are associated with actively transcribed regions in the human genome”. en. In: DNA Res. 23.5 (Oct. 2016), pp. 477–486.

[9] Tae-Young Roh. “ChIP-Seq strategy to identify Z-DNA-forming sequences in the human genome”. en. In: Methods Mol. Biol. 2651 (2023), pp. 167–177.

[10] Ryan S Czarny and P Shing Ho. “Thermogenomic analysis of left-handed Z-DNA propensities in genomes”. en. In: Methods Mol. Biol. 2651 (2023), pp. 195–215.

[11] Guangchuang Yu and Qing-Yu He. “ReactomePA: an R/Bioconductor package for reactome pathway analysis and visualization”. en. In: Mol. Biosyst. 12.2 (Feb. 2016), pp. 477–479.

[12] Fidel Ramírez et al. “deepTools2: a next generation web server for deep-sequencing data analysis”. en. In: Nucleic Acids Res. 44.W1 (July 2016), W160–W165.

[13] Hadley Wickham. Ggplot2. en. 1st ed. Use R! New York, NY: Springer, Dec. 2009.

[14] Johan Larsson. eulerr: Area-Proportional Euler and Venn Diagrams with Ellipses. R package version 7.0.2. 2024. url: https://CRAN.R-project.org/package=eulerr.

[15] Niklas Bruse and Simon J van Heeringen. “GimmeMotifs: an analysis framework for transcription factor motif analysis”. Nov. 2018.

[16] Simon J van Heeringen and Gert Jan C Veenstra. “GimmeMotifs: a de novo motif prediction pipeline for ChIP-sequencing experiments”. en. In: Bioinformatics 27.2 (Jan. 2011), pp. 270–271.

[17] Raquel B Haga and Anne J Ridley. “Rho GTPases: Regulation and roles in cancer cell biology”. en. In: Small GTPases 7.4 (Oct. 2016), pp. 207–221.

[18] Shuchen Chen et al. “Expression and prognostic analysis of Rho GTPase-activating protein 11A in lung adenocarcinoma”. en. In: Ann. Transl. Med. 9.10 (May 2021), p. 872.

[19] Sung Chul Ha et al. “The crystal structure of the second Z-DNA binding domain of human DAI (ZBP1) in complex with Z-DNA reveals an unusual binding mode to Z-DNA”. en. In: Proc. Natl. Acad. Sci. U. S. A. 105.52 (Dec. 2008), pp. 20671–20676.

